# Hedgehog pathway inhibitors significantly reduce the formation of heterotopic ossification in a direct trauma/burn mouse model

**DOI:** 10.1101/2021.01.31.429058

**Authors:** Atanu Chakraborty, Jelena Gvozdenovic-Jeremic, Fang Wang, Stephen W. Hoag, Ekaterina Vert-Wong, Ryan M. Pearson

**Author notes:** These authors contributed equally to this study (Atanu Chakraborty, Jelena Gvozdenovic-Jeremic).

## Abstract

Heterotopic ossification (HO), either acquired or hereditary, is featured by ectopic bone formation outside of the normal skeleton. The acquired form of HO is a debilitating and common complication of musculoskeletal trauma, central nervous system injury, burns, combat trauma, hip and elbow fractures, and total joint replacement surgeries. It can be characterized as abnormal bone formation that occurs mostly by endochondral ossification. Recent studies have implicated inflammation and dysregulation of Hedgehog (Hh) signaling as major early contributors to HO formation. Here, we demonstrate that administration of the Hh pathway inhibitor, arsenic trioxide (ATO), prevented acquired HO in a clinically-relevant trauma/burn mouse model. We further evaluated the effects of two additional Hh pathway antagonists: cholecalciferol and pravastatin on mitigating osteoblast differentiation. Finally, we assessed the effect of a combination of Hh pathway inhibitors on reducing systemic proinflammatory responses. A targeted combination approach using Hh pathway inhibitors may offer potential therapeutic benefits though targeting differential components of the Hh pathway. Taken together, our study demonstrates that the administration of single or multiple Hh pathway inhibitors may have the potential to reduce the formation of acquired HO.

## 1. Introduction

Heterotopic ossification (HO) refers to lamellar bone formation inside soft tissue, where bone should not exist [1]. HO is a pathology triggered by injury or mutations for which currently there is no definite treatment. There are two forms of HO: genetic and acquired. Genetic mutations in the type 1 bone morphogenetic protein (BMP) receptor, ALK2, cause the rare hereditary form of HO, such as Fibrodysplasia Ossificans Progressiva (FOP) [2, 3]. Acquired HO is more common and triggered by traumatic injury seen following fractures, dislocations, burns, or after certain operative procedures such as total hip arthroplasty [4], open treatment of elbow fractures [5], and after intramedullary fixation of femoral shaft fractures [6]. Both, FOP and acquired HO are formed primarily by endochondral ossification [7, 8]. Endochondral ossification is a natural development process of long bone as well as fracture healing, which begins with mesenchymal stem cell (MSC) differentiation and chondrogenesis as opposed to intramembranous ossification that involves direct differentiation of MSCs into osteoblasts [9]. Although the molecular and cellular mechanisms that lead to HO remain largely unknown, the inappropriate differentiation of MSCs induced by a pathological imbalance of local or systemic factors (such as inflammation) contribute to HO formation [10].

Previous studies have demonstrated that activation of the Hedgehog (Hh) signaling pathway is sufficient and necessary for HO formation through the intramembranous ossification mechanism, suggesting that this signaling crosstalk may also be important in physiological bone formation and homeostasis [11]. Furthermore, a pathological link between the Hh signaling pathway and endochondral HO has been recently postulated [12]. Specifically, it was shown that local dysregulation (activation) of Hh signaling participates in HO’s pathophysiology of murine muscle injury-induced HO model (Nse-BMP4) [13].

Acute proinflammatory responses have a proven role in the development of HO [10]. Following injury, an inflammatory cascade is initiated by cells of the innate and adaptive immune system such as neutrophils, macrophages, B cells, and T cells. Proinflammatory markers such as monocyte chemoattractant protein-1 (MCP-1) and interleukin-6 (IL-6), and several others are secreted to attract circulating immune cells to the injury and act to modulate the local environment [14]. The core mechanisms of HO development are multiple disease-specific cellular and extracellular molecular elements that form a unique local microenvironment, particularly those of injury-induced stem cell niche, which regulates the proliferation and osteogenic differentiation of MSCs. Infiltrating MSCs can differentiate into bone cells under inflammatory conditions [15, 16]. Inflammatory cytokines induce osteogenic MSC gene expression and differentiation by enhancing runt-related gene 2 (*RUNX2*) expression and alkaline phosphatase (*ALP*) gene expression [17]. This phenomenon indicates the involvement of MSCs in bone homeostasis in the context of inflammation induced by trauma.

The burn/tenotomy model recapitulates numerous features associated with endochondral ossification, which is driven in part by Hh signaling, and the primary form of HO induced in patients following traumatic injury. Additionally, this model accurately reproduces the inflammatory states that cause the majority of cases of HO. The model protocol involves creating a 30% total body surface area partial thickness contact burn on the dorsal skin as well as division of the Achilles tendon at its midpoint. Relying solely on traumatic injury to induce HO at a predictable location allows for the time-course study of endochondral heterotopic bone formation from intrinsic physiologic processes and environment only [18, 19].

In this study, we demonstrate the role of single or combinations of Hh pathway inhibitors as a potential therapeutic strategy for acquired HO. Our recent *in vitro* study showed Hh pathway inhibitors prevented early osteogenesis of bone marrow MSCs [20]. Using the burn/tenotomy mouse model of HO [18], we evaluated the effect of arsenic trioxide (ATO) on HO formation. Arsenic trioxide (ATO) is a potent Hh pathway inhibitor which binds directly to the GLI1 transcriptional factor [21]. ATO is among the first-line chemotherapeutics used in oncological practice. It has shown substantial efficacy in treating patients with relapsed or refractory acute promyelocytic leukemia [22]. Next, we evaluated two additional Hh inhibitors, cholecalciferol, and pravastatin and their impact on mineralization and osteogenic gene expression. Previously, exogenous addition of cholecalciferol has been shown to inhibit the Hh pathway in a basal cell carcinoma (BCC) cell line by efficiently inhibiting Smo from translocating into the primary cilium [23]. Statins, 3-hydroxy-3-methylglutaryl (HMG)-CoA reductase inhibitors, block cholesterol synthesis which is required for Hh signaling transduction [24]. Cholesterol was recently investigated for its role in MSC osteogenic differentiation where endogenous cholesterol was required for Hh signaling induced osteoblast differentiation [25]. Finally, we demonstrate the effect of a combination Hh pathway inhibitor therapy on the systemic proinflammatory response associated with HO. Our results indicate that the application of single or multiple Hh pathway inhibitors has the potential to significantly reduce the formation of acquired HO.

## 2. Materials and Methods

### 2.1. Materials

Arsenic trioxide (ATO): Assay ≥ 99.9%, Millipore Sigma (St. Louis, MO); Cholecalciferol (Vitamin D3): HPLC, Assay ≥ 98%, Pravastatin Na: HPLC, Assay ≥ 98.8%, Tocris (Minneapolis, MN). Sterile Water for Irrigation (USP; Frederick, MD), Baxter (Round Lake, IL). Polyoxyl 35 castor oil (USP-NF; Frederick, MD), Kolliphor^®^ EL (BASF; Jessup, MD). Tween-80, PEG-400, neutral buffered formalin were purchased Millipore Sigma (St. Louis, MO). Aluminum 6061 35 g weight (2 cm x 2 cm x 3 cm) was purchased from eMachineShop^®^ (Mahwah, NJ). 5-0 vicryl sutures were purchased from Fisher Scientific (Waltham, MA). Tegaderm dressings were purchased from VWR (Radnor, PA). Buprenorphine was obtained from MWI Animal Health (Boise, ID).

### 2.2. OsteoImage Mineralization Assay

The OsteoImage Mineralization assay was performed following the manufacturer’s instructions (Lonza, Walkersville, Inc). This Assay can quantify *in vitro* mineralization by osteogenic stem cells, primary osteoblasts, and osteoblast-like cell lines. Osteoblast differentiation is marked by the formation of mineralized nodules composed of inorganic hydroxyapatite (HA) (Ca_10_(PO_4_)_6_(OH)_2_) and organic components including type I collagen. The OsteoImage^™^ Assay is based on the specific binding of the fluorescent OsteoImage^™^ Staining Reagent to the hydroxyapatite portion of the bone-like nodules deposited by cells. Human bone marrow stem cells (hBMSCs) were plated in triplicates in 96 well plates. hBMSC were obtained from ATCC (ATCC PCS 500-012, Manassas, Virginia, USA) and propagated in basal medium (ATCC PCS 500-030) supplemented with ATCC growth kit (ATCC PCS 500-041) in a humidified atmosphere with 5% CO_2_ at 37°C. For hBMSCs passage 1-3 was used for all experiments. After the cells reached confluence, the medium was replaced with osteogenic differentiation medium (OM). OM consisted of: Dulbecco’s Modified Eagle Medium (DMEM), 10% fetal bovine serum (FBS), 100 U/mL penicillin, 100 μg/mL streptomycin, 2 mM glutamine (Gibco; Waltham, MA), 10 μM L-ascorbic acid 2-phosphate (Wako Chemicals; Richmond, VA) and 10 mM β-glycerol phosphate (Millipore Sigma (St. Louis, MO). Triplicate wells were exposed to differentiation media without any agent (appropriate vehicle only-control), or with cholecalciferol or pravastatin for 21 days. Media replacement occurred every 2-3 days with fresh OM (made each time during feeding) with or without single agents. The results were obtained using a fluorescence microplate reader with 492/520 excitation/emission wavelength.

### 2.3. RT–PCR

We isolated the total RNA first with Trizol (Invitrogen Corporation; Carlsbad, CA) and then with RNeasy Kit (Qiagen, Germantown, MD). cDNA was generated using the High Capacity cDNA synthesis kit (Applied Biosystems; Foster City, CA). Quantitative real-time (RT)-PCR was performed using a Bio-Rad cycler (Hercules, CA) and 40 cycles of 95°C for 15 seconds and 60°C for 60 seconds. PCR product was detected using SybrGreen (Bio-Rad; Hercules, CA). Primers for human BMSC cells used for amplifications are: *GAPDH* Forward: 5’-GGC ATG GAC TGT GGT CAT GAG-3’; Reverse: 5’-TGC ACC ACC AAC TGC TTA GC-3’ *ALP*: Forward 5’-CTC CCA GTC TCA TCT CCT-3’, Reverse 5;-AAG ACC TCA ACT CCC CTG AA; *RUNX2* Forward: 5’-ACT TCC TGT GCT CGG TGC T-3’; Reverse 5’-GAC GGT TAT GGT CAA GGT GAA-3’.

### 2.4. Preparation of Hedgehog Pathway Inhibitor Solutions

Stock solutions of ATO (1 mg/mL), pravastatin sodium (20 mg/mL) and cholecalciferol (vitamin D3) (0.5 mg/mL) were prepared using Sterile Water for Irrigation. In the vitamin D3 stock solution, Kolliphor^®^ EL was added at a concentration was 1 mg/mL as a solubilizing agent, i.e., the vitamin D3 and Kolliphor^®^ EL were present at a fixed ratio. Based on the target formulation in **Table 1**, a measured volume of each stock solution was added to a 20 mL amber bottle, capped, and hand shaken until uniformly mixed. For each formulation, 10 mL was produced and then transferred by syringe into 2 mL sterile amber vials labeled with the corresponding identification number.

**Table 1.**
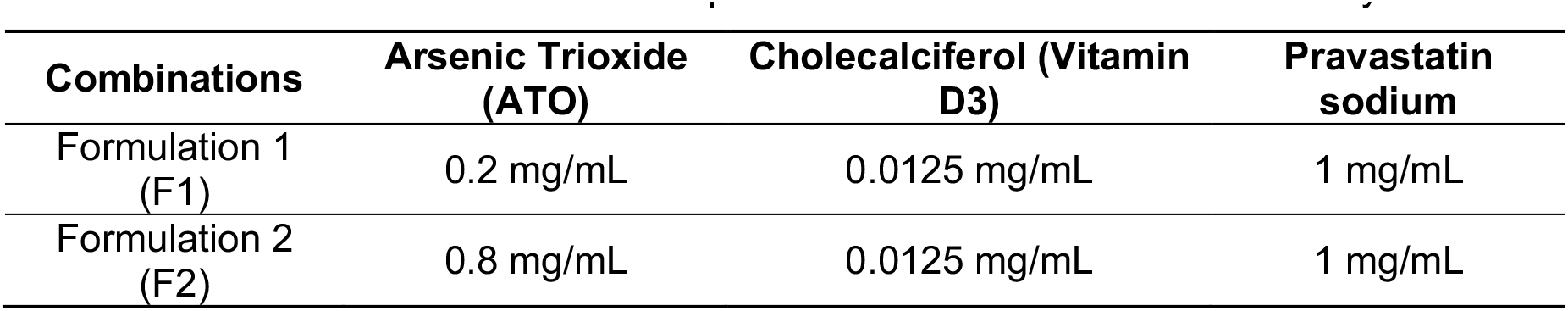
Combinations of multiple Hh inhibitors evaluated in this study.

### 2.5. In vitro Osteogenesis

hBMSCs were plated in triplicates in 96 well plates and cultured as described above. Triplicate wells were exposed to OM without any formulation (Kolliphor^®^ EL-control), or with F1 or F2 diluted 500x in OM for six days. Media replacement occurred every other day with fresh OM (made each time during feeding) with or without Kolliphor^®^ EL or F1 or F2 (500x dilution). At day six cells were harvested for RT-PCR analysis.

### 2.6. Mice

Male C57BL/6J (5-7 weeks old) were purchased from The Jackson Laboratories (Bar Harbor, ME). The mice were housed under specific pathogen-free conditions at the University of Maryland, Baltimore. All mice procedures and experiments were compliant to the protocols of the University of Maryland, Baltimore Animal Care and Use Committee (IACUC: #0818014).

### 2.7. Induction of HO

Ectopic bone formation was induced as previously described using the burn/tenotomy mouse model [18]. Briefly, mice were anesthetized using inhaled isoflurane. The dorsum of the mouse as well as the hind paw were shaved to accommodate the burn injury and allow visualization of the surgical site. An Achilles’ tenotomy was performed by dissection of the midpoint of the left Achilles’ tendon followed by closing the wound with a single 5-0 vicryl stitch. To induce the 30% total body surface area burn, a 35 g aluminum weight heated to 60°C was applied to the shaved dorsum for 18 seconds continuously. The burn site was dried and a tegaderm dressing was applied. Post-operative pain was managed with subcutaneous 0.01 mg/kg buprenorphine injections every 6-8 hours for 3 days. Immediately post-surgery, mice were administered saline (negative control), Kolliphor^®^ EL vehicle (selected candidates’ negative control), or ATO (2 or 5 mg/kg) by intraperitoneal injection. The mice were dosed every day for the duration of the study unless toxicity is observed. For the detection of serum biomarkers study, mice were administered the respective negative control, F1, or F2 every 2-3 days for one week (3 injections total).

### 2.8. Detection of serum biomarkers for HO formation

Mice were anesthetized at 1-week post-HO induction and after receiving three doses of F1 and F2 formulations. Blood was collected by cardiac puncture and serum was separated from whole blood by centrifugation for 5 min at 600 x g. Enzyme-linked immunosorbent assays (ELISA) (BioLegend; San Diego, CA) were used to measure murine interleukin-6 (IL-6) and monocyte chemotactic protein-1 (MCP-1).

### 2.9. Micro-CT analysis of ectopic bone formation

To assess the effects of ATO treatment on ectopic bone formation, we used micro-CT imaging to quantify the reduction in total and soft tissue bone volumes. Images were acquired using a Siemens Inveon small animal PET-CT at the Core for Translational Research in Imaging (CTRIM) at the University of Maryland, Baltimore. Image acquisition and post-acquisition data analysis using the Inveon Research Workspace. Acquired images were fused and the total HO volume was calculated for the control and ATO treatment groups.

### 2.10. Histological evaluation of HO formation

The alteration in architecture of HO formation by ATO was assessed at 9 weeks post-HO induction by histology. Hind limbs were formaldehyde fixed and subsequently decalcified for three weeks using an 10% EDTA solution. Hind limbs were then paraffin embedded and sectioned in 5-7 μm slices. Slides were stained for H&E to define histological architecture and Masson’s Trichrome staining using standard procedures by the Pathology Biorepository Shared Services Core at University of Maryland, Baltimore [26, 27].

## 3. Results

### 3.1. ATO reduces HO in a dose dependent manner in burn/tenotomy HO mouse model

We quantified both, calcaneal and soft tissue HO formation at 9 weeks post-surgery using micro-CT imaging. The burn/tenotomy model allows for predictable ectopic bone formation along the calcaneus as well as proximally within the soft tissue. Soft tissue HO specifically forms within the proximal transected tendon and distal gastrocnemius but away from the calcaneus [28]. Two groups of ATO were evaluated at increasing doses of 2 mg/kg (ATO-L (low)) and 5 mg/kg (ATO-H (high)). ATO was administered daily by intraperitoneal injection. The control for this experiment was saline (No Tx). Quantitative measurement of the bone volume showed that ATO-H significantly reduced the HO in both, soft tissue and calcaneal area around the tendon (**Figure 1**) (n=5-6). The reduction in bone volume corresponded to a decrease of 50% compared to both No Tx and ATO-L groups.

**Figure 1.**
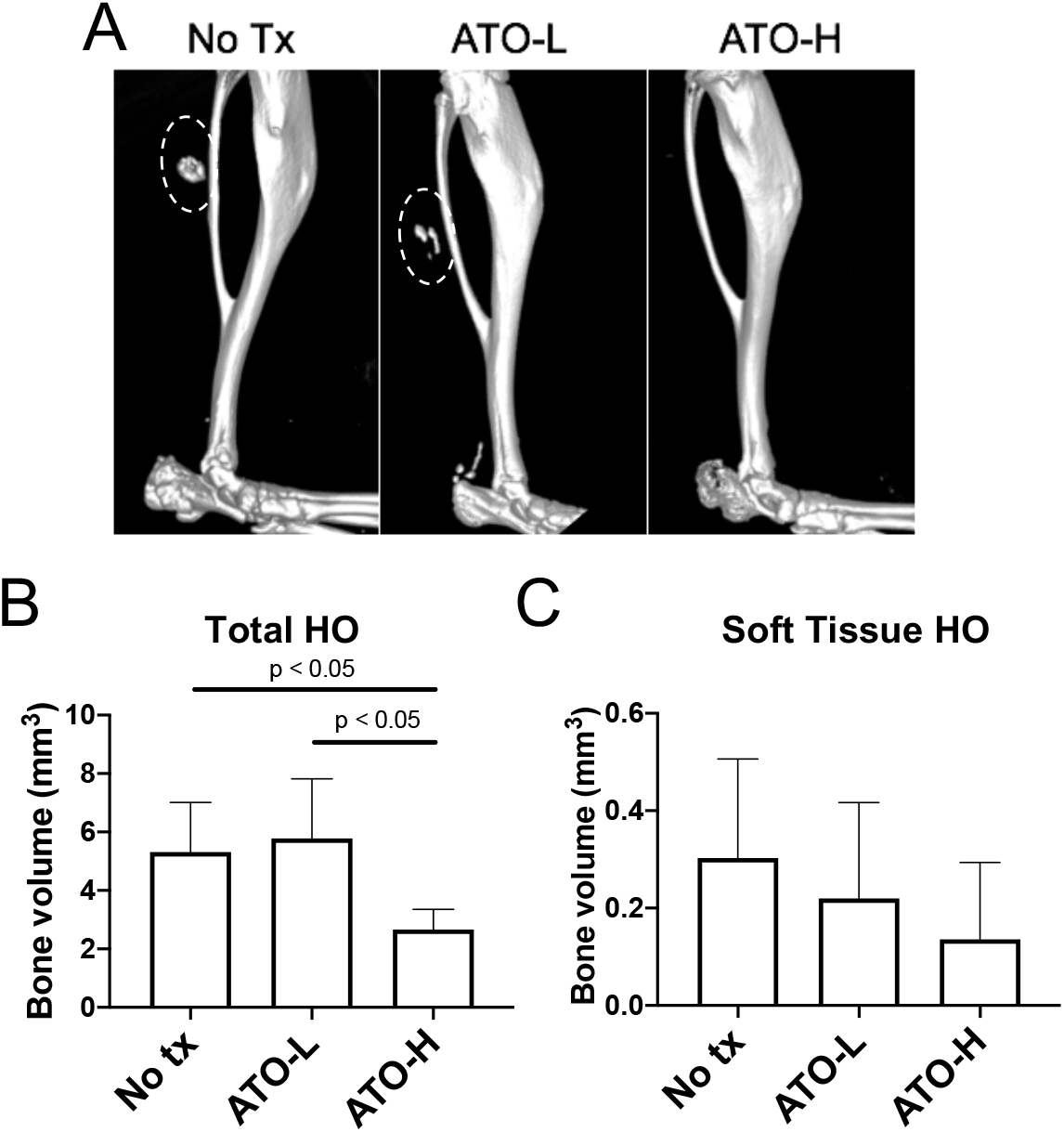
C57BL/6 mice were treated every day 100 μL (ATO-L) and 250 μL (ATO-H) *via* injection intraperitoneally (ATO 0.5 mg/mL), or normal saline (No Tx) for 9 weeks. (A) Three-dimensional reconstruction of micro-CT scans of ATO and control-treated burn/tenotomy mice at 9 weeks post-injury. *i*) No tx (saline); *ii*) ATO-L; *iii*) ATO-H; (B) Quantification of total HO; (C) Quantification of soft tissue HO. Dotted circles denote the location of soft tissue HO. Statistical differences between groups were determined by performing a one-way ANOVA and Tukey’s post hoc test. Error bars (S.D). n=5-6 per group.

### 3.2. Histological changes in calcaneal cross sections

The histology of *in vivo* bone formation was examined by H&E and Masson’s trichrome staining. Representative histological sections are depicted in **Figures 2 and 3**. Brightly red regions in H&E sections represent the newly formed heterotopic bone areas and were observed in No Tx (control) and ATO-L groups [26]. Importantly, newly formed HO was not observed in the ATO-H group (**Figure 2**). These findings were further corroborated by Masson’s trichrome staining, where newly formed immature bone (purple regions) was not present in samples obtained from the mice treated with high dose ATO (**Figure 3**). Newly formed bone regions, or purple color, indicates that the collagen fibers, or osteoids, are present, while the absence of bright red color indicates the bone is not fully mature yet.

**Figure 2.**
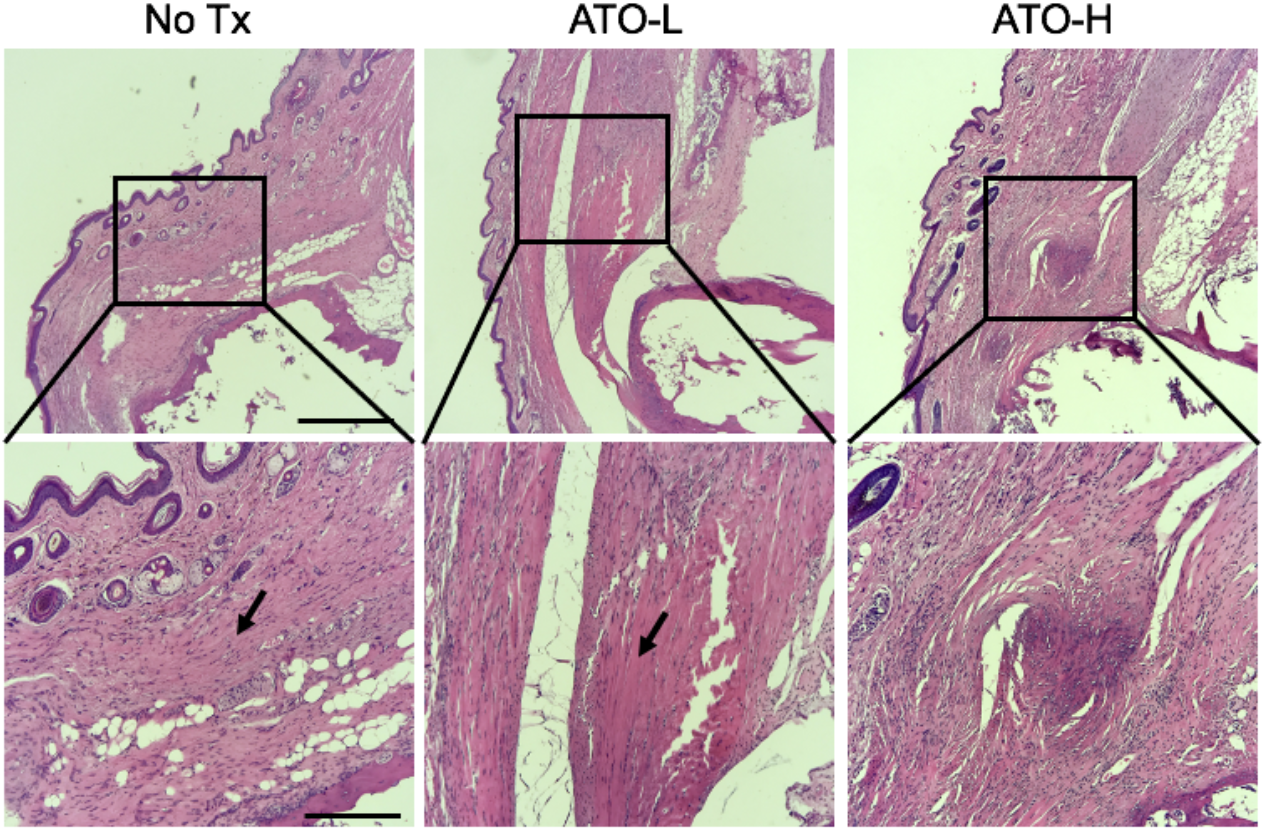
Histological images of representative calcaneal cross sections of various treatment groups at 9 weeks post injury (H&E stain, ×40, ×100). Arrowheads point to the newly formed bone. No Tx, saline treatment; ATO-L (2 mg/kg treatment); ATO-H (5 mg/kg treatment). Scale bar (550 μm: top; 220 μm: bottom).

**Figure 3.**
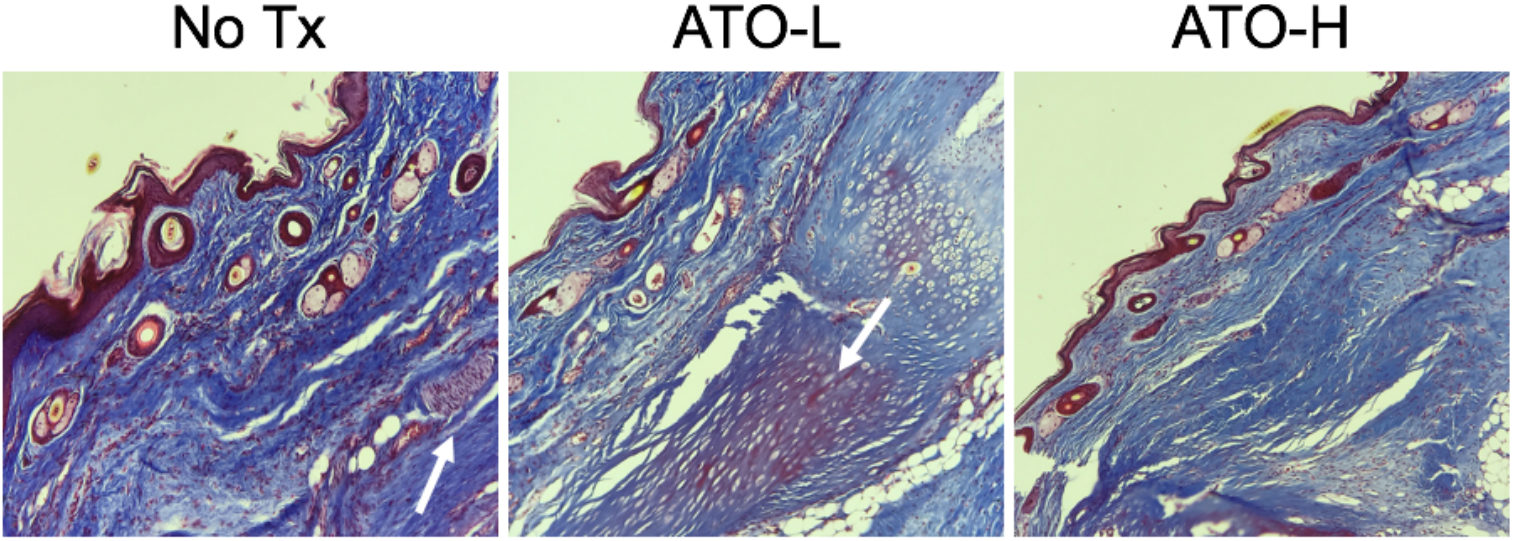
Masson’s trichrome images of representative calcaneal cross sections of various treatment groups. Timepoint 9 weeks. Magnification 100X. Purple stain (arrows) represents area of newly formed immature bone. No Tx, saline treatment; ATO-L (2 mg/kg treatment); ATO-H (5 mg/kg treatment).

### 3.3. Cholecalciferol and Pravastatin block in vitro mineralization of hBMSC

Exogenous addition of cholecalciferol has been shown to inhibit the Hh pathway in a basal cell carcinoma (BCC) cell line [23, 29, 30]. Interestingly, studies have found that cholesterol synthesis is required for Sonic Hh (SHH) signaling [31, 32]. Thus, blocking cholesterol synthesis with statins might be beneficial for HO mitigation. We further evaluated these two Hh antagonists: cholecalciferol and pravastatin for inhibiting osteoblast differentiation of hBMSCs using an *in vitro* OsteoImage mineralization assay. The cells were supplemented with OM for 21 days, with or without (control) Hh antagonist. As shown in **Figure 4**, both cholecalciferol and pravastatin blocked mineralization of hBMSCs at all concentrations evaluated (0.1 μM to 0.5 μM) at 21 days. These results suggest that sterols might be a new therapeutic target for the treatment of HO, and readily available agents such as statins (HMG-CoA reductase inhibitors) or vitamin D might offer more safe and efficacious ways of prevention and treatment of HO.

**Figure 4.**
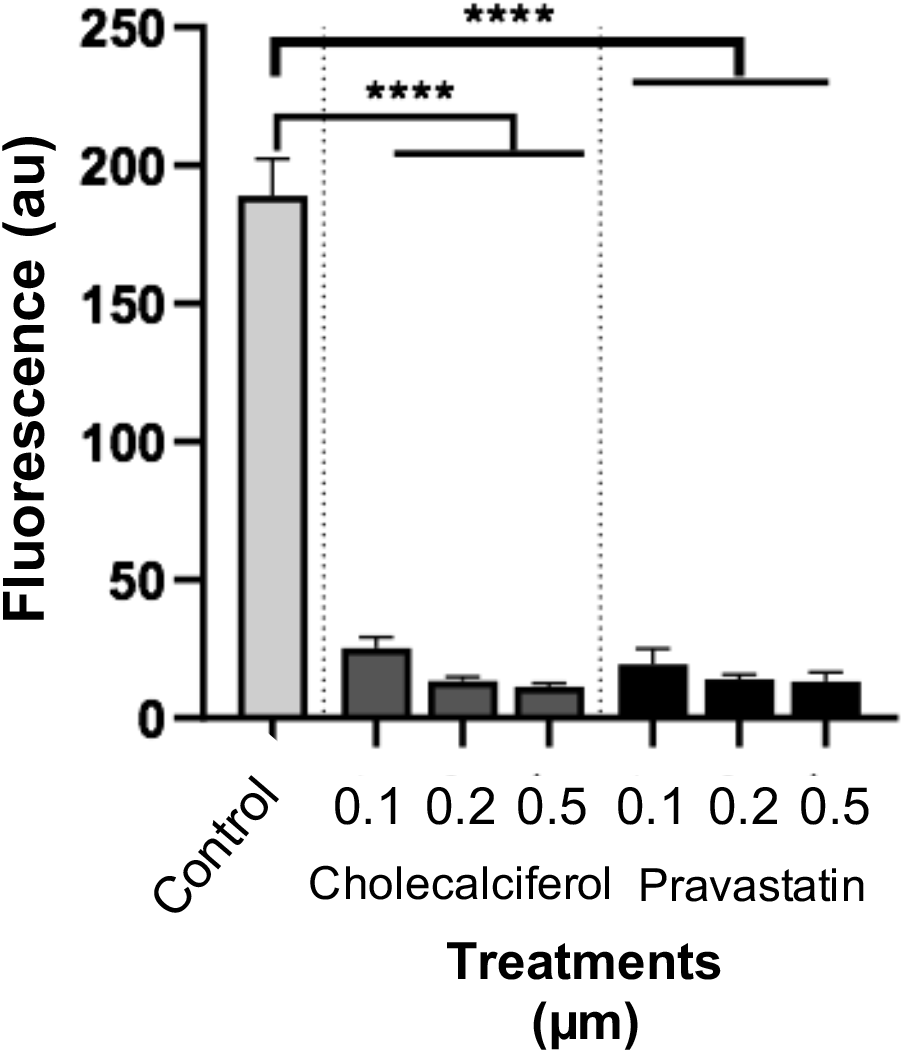
Mineralization assay of hBMSC cultured with osteo-differentiation factors. Mineralization evaluated by staining cells using an OsteoImage Assay. Graphs depict relative fluorescence of hBMSC during osteogenesis 21 days post-treatment with Cholecalciferol (0.1-0.5 μM) and Pravastatin (0.1-0.5 μM) compared to the control (no treatment). Statistical differences between groups were determined by performing a one-way ANOVA and Tukey’s post hoc test. Error bars (SEM). n=3 per group.

### 3.4. In vitro combination therapy testing

In line with the *in vitro* and *in vivo* single agent Hh inhibitor data, we developed two combinations of three Hh inhibitors (ATO, pravastatin, cholecalciferol) as described in **Table 1** that act on three distinct Hh pathway checkpoints [33–35]. These formulations were tested *in vitro* to investigate their effects on inhibiting an early osteogenesis, revealed by *ALP* and *RUNX2* expression. Each formulation was found to significantly suppress the differentiation of mesenchymal progenitor stem cells into osteoblasts (**Figure 5**). The formulations exhibited a significant decrease from 25% to 50% respectively in alkaline phosphatase (*ALP*) mRNA levels compared to control (Kolliphor^®^ EL). Comparable results were also obtained with quantification of *RUNX2*, an early osteoblast formation gene marker. The advantage of using this multicomponent approach lies in its unique ability to target different checkpoints of the Hh signaling pathway involved in early HO formation, thereby efficiently inhibiting trans-differentiation of MSCs into osteoblasts.

**Figure 5.**
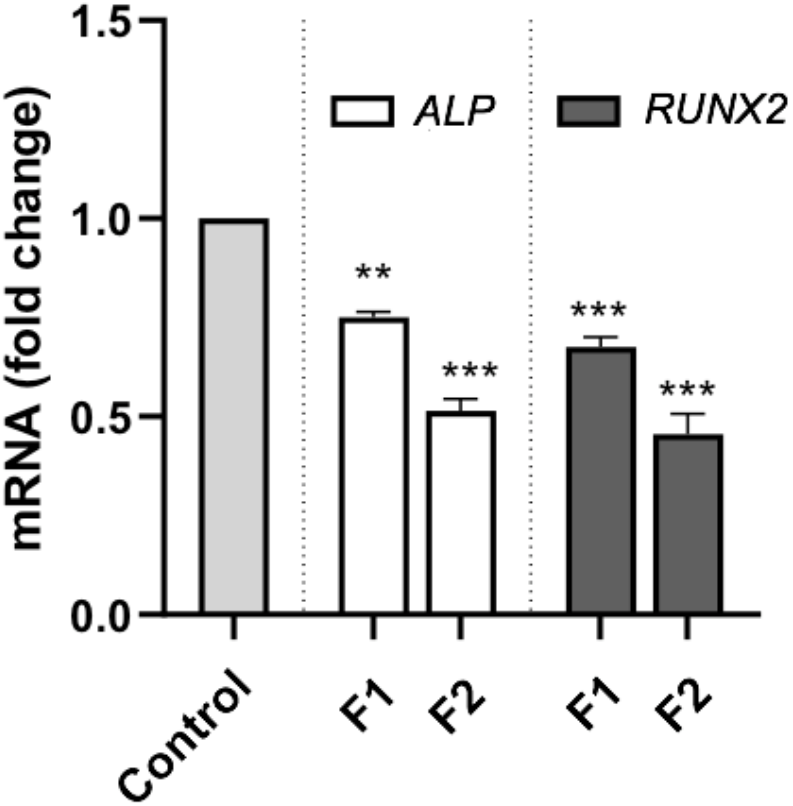
*In vitro* combinatorial formulation analyses of Hh inhibitors. Graphs depict relative gene expression of Alkaline Phosphatase (*ALP*) and runt-related gene 2 (*RUNX2*) genes of hBMSC during osteogenesis 6 days post-treatment with formulations F1 and F2 (diluted 500x). F1 (ATO: 0.2 mg/mL; Pravastatin: 1 mg/mL; Vitamin D3: 0.0125 mg/mL). F2 (ATO: 0.8 mg/mL; Pravastatin: 1 mg/mL; Vitamin D3: 0.0125 mg/mL). Control is placebo, Kolliphor^®^ EL (1 mg/mL) (diluted 500x). Statistical differences between groups were determined by performing a one-way ANOVA and Dunnett’s post hoc test. Error bars (SEM). n=3 per group.

### 3.5. Hh-inhibitor combinations elicit an anti-inflammatory response in burn/tenotomy HO mouse model

A systemic inflammatory state is associated with the development of HO [10, 14]. Studies have shown a prognostic correlation between inflammatory cytokine concentrations and HO development [36, 37]. It has been previously demonstrated that inflammatory cytokine concentrations in the serum are maximally between 2 days and 1 week-post tenotomy and decreased to week 3 [36]. To understand the effects of formulations F1 and F2 on the systemic inflammatory response, we assessed inflammatory cytokine secretions in the serum at 1 week post-HO induction, after receiving three doses of respective formulations. Mice were terminally bled by cardiac puncture and serum was separated from whole blood. ELISA was used to measure monocyte chemoattractant protein-1 (MCP-1) and interleukin-6 (IL-6) (**Figure 6**). Both F1 and F2 significantly reduced the serum levels of MCP-1 compared to the placebo (Kolliphor^®^ EL-containing vehicle). Interestingly, we found that F2 was more effective than F1, which was attributed to the higher concentration of ATO present in the formulation. No significant differences were observed related to IL-6 expression, however, F2 did trend towards significance (p = 0.07). The results obtained in this experiment indicated the potential for a combination therapy of Hh inhibitors to modulate inflammatory responses associated with traumatic injury-associated HO. Future studies are planned to systematically evaluate the role of ATO concentration alone or in combination with other Hh inhibitors to identify potential synergistic opportunities for HO treatment development.

**Figure 6.**
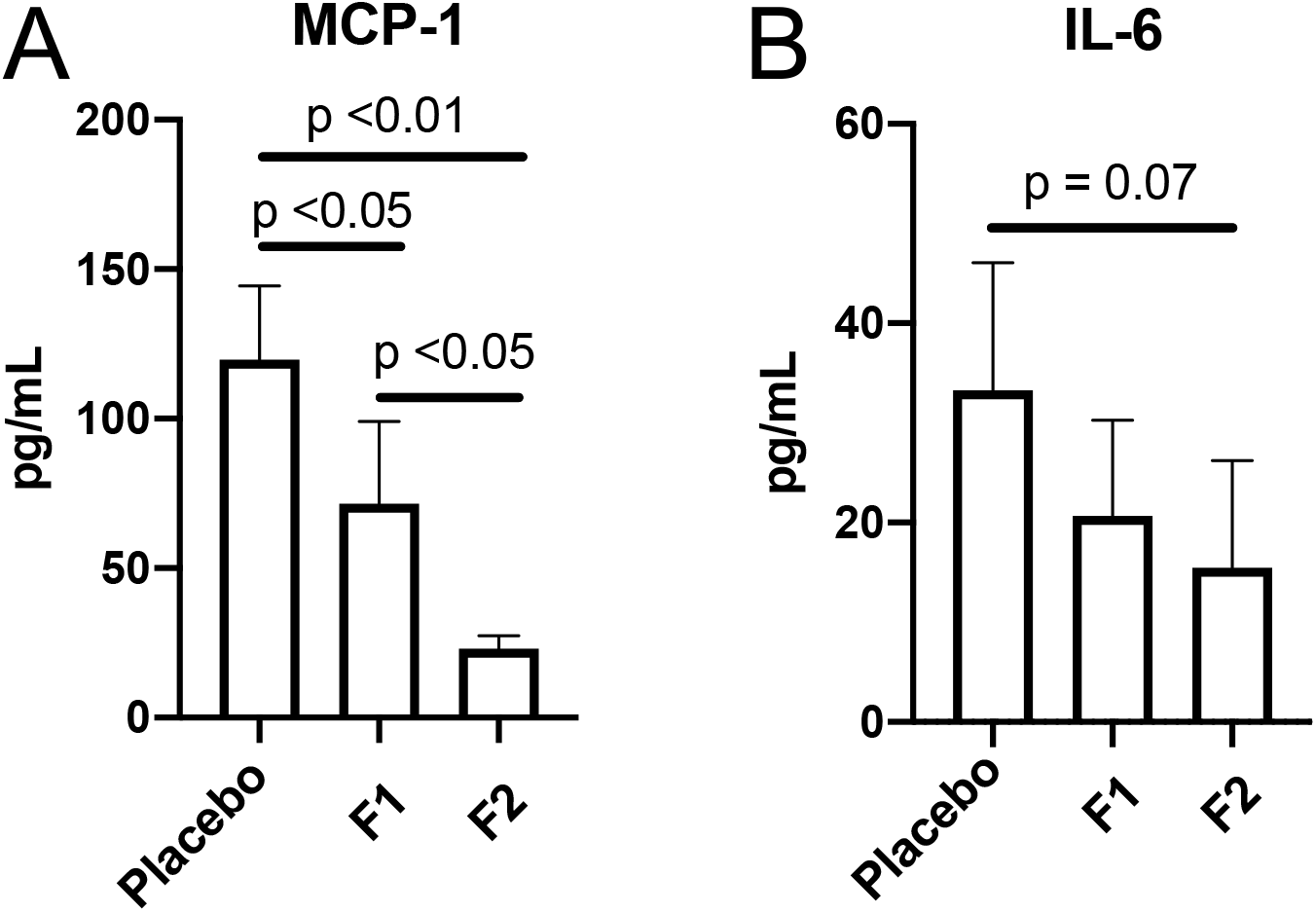
Analysis of serum cytokine levels in the burn/tenotomy model of HO. Graphs depict concentrations of monocyte chemoattractant protein-1 (MCP-1) (A) and interleukin-6 (IL-6) (B) in serum from mice 7 days post-treatment. Mice were treated with a total of 3 doses (50 μL per injection) of respective formulations. Statistical differences between groups were determined by performing a one-way ANOVA and Tukey’s post hoc test. Error bars (S.D). n=4-5 per group.

## 4. Discussion

The primary risk factor for acquired HO is inciting trauma or flare-ups in FOP. Additionally, inflammation has been shown to play a significant role [38]. Numerous mouse models have been developed to investigate the molecular mechanisms of HO formation as well as to evaluate potential treatments [16]. However, these models rely on the administration of exogenous substances, exogenous cell constructs, or genetic mutations. The direct trauma/burn model is an acquired model of HO, where HO is precipitated by trauma (burn) or has a neurogenic cause [39]. Inducing HO by direct trauma/burn accurately and precisely enables the efficacy of any therapeutic agent to be investigated due to the formation of HO in predicable locations and within a known time-course [18]. Like HO observed in the clinic, the reactive bone in this model can adjoin with skeletal bone or develop remotely in soft tissue. Soft tissue HO forms *de novo*, without the influence of adjacent cartilage, bone or periosteum typically located in close proximity to extraskeletal bone at the calcaneus.

The Hh pathway has been shown to be indispensable in osteogenesis [40, 41]. The effectiveness of Hh pathway inhibitors for mesenchymal progenitor cells differentiation was demonstrated in case of rare genetic disease-progressive osseous heteroplasia (POH) [11]. In POH, a human disease caused by null mutations in GNAS, which encodes Gαs, Hh signaling is upregulated in ectopic osteoblasts and HO is formed by intramembranous ossification. To the best of our knowledge, prevention of HO using Hh pathway antagonists in an acquired HO mouse model has not been reported. Although, Kan et al. identified the Hh pathway as an important contributing factor in newly formed HO lesions, they found that global inhibition of Hh signaling was insufficient to block HO in Nse-BMP4 transgenic mouse model of acquired HO [12]. Indeed, the lower dose of ATO intraperitoneal administration at 2 mg/kg was also ineffective in our study and burn/tenotomy mouse model (**Figure 1**). Increasing the dose of ATO to 5 mg/kg significantly reduced the newly formed bone volume by 50%. As our data suggest, the dosing regimen and the frequency of application in systemic administration is crucial to achieve effectiveness of Hh inhibitor treatment. We hypothesize that dosing and frequency are critical for reaching the effective local concentration of the Hh inhibitor at the injury. However, systemic administration of high ATO doses might pose a general concern due to toxicity [42]. In particular, high dose administration of ATO (10 mg/kg) to Sprague-Dawley rats resulted in a reduction in trabecular bone volume (femur) but not 5 mg/kg [43], the same dose of ATO used in our present study. During the course of our study, we did not observe lethargy or weight loss due to Hh pathway inhibitor administration in mice during the course of the treatment (data not shown), suggesting that resorption of normal endochondral bone did not occur. Histological examination of the calcaneal region was performed to identify alterations in tissue and bone architecture (**Figures 2 and 3**). Our work has demonstrated that administration of an Hh inhibitor (ATO) effectively reduced HO formation and highlights their potential therapeutic use for treating the acquired form of HO.

Our proposed combination approach has the potential to target critical checkpoints in the Hh signaling pathway, which may significantly improve safety and efficacy of the Hh pathway inhibitors by overcoming redundancies in Hh signaling and therefore drug resistance [44]. A synergistic interaction has the potential to allow the application of lower doses of the combination constituents, a situation that may reduce adverse reactions. There are examples in the literature where combination of Hh pathway inhibitors produce more potent inhibition at lower drug doses [45, 46]. The IC50 of ATO with a combination of an upstream Smo inhibitor can be reduced up to 12-fold [46]. Our recently published data also demonstrate the potential use of statins and cholecalciferol as therapeutic agents in a combination for the HO treatment [20]. Here, we have developed two formulations and found *in vitro* that they elicit significant potential of suppressing the differentiation of progenitor stem cells into bone cells (**Table 1** and **Figure 5**). Additionally, these formulations significantly reduced serum proinflammatory cytokine levels, thereby suppressing initial inflammatory responses that play a contributing role in MSC differentiation and subsequent HO formation. Administration of F2 resulted in an 81% decrease for MCP-1 and 54% for IL-6 compared the placebo control (**Figure 6**). Given the demonstrated importance of suppressing systemic inflammatory responses in addition to the Hh signaling pathway to mitigate HO, our results provide a strong rationale for future studies aimed at identifying and evaluating synergistic combinations of Hh pathway inhibitors.

## 5. Conclusions

Development of new drugs for HO has been hindered due to a complex etiology and underlying pathophysiology. Our work has demonstrated that ATO, a Hh pathway inhibitor, effectively and safely reduced HO formation using an acquired HO mouse model, which could implicate its therapeutic potential for endochondral HO formation, such as FOP. Two additional Hh pathway inhibitors, cholecalciferol and pravastatin, were shown to reduce mineralization and osteogenic gene expression of hBMSCs. Finally, a combination of all three Hh pathway inhibitors resulted in significant reductions in systemic proinflammatory responses. Taken together, our findings using multiple Hh pathway inhibitors represent a potential therapeutic advancement for the treatment of acquired HO.

## Author Contributions

Conceptualization, JGJ, EWV, SWH and RMP.; methodology, AC, JGJ, and FW; validation, AC, FW, SWH and RMP; formal analysis, AC, RMP, EWV, JGJ and SWH.; resources, SWH and RMP; data curation, AC and FW.; writing—original draft preparation, AC and JGJ; writing - reviewing & editing, AC, JGJ, EVW, and RMP; supervision, SWH, EVW, and RMP.

## Acknowledgments

We thank Dr. David Fink from bwtech@UMBC Research & Technology Park at the University of Maryland, Baltimore County for his helpful discussions regarding experimental planning. We thank the University of Maryland School of Medicine Center for Innovative Biomedical Resources, Translational Research in Imaging @ Maryland (CTRIM) – Baltimore, Maryland for their help with acquiring and processing the micro-CT imaging. We also thank the University of Maryland School of Medicine Center for Innovative Biomedical Resources, Pathology Biorepository Shared Services Core – Baltimore, Maryland for their help preparing the H&E and Masson’s trichrome histology sections.

